# The dynamics of resting fluctuations in the brain: metastability and its dynamical cortical core

**DOI:** 10.1101/065284

**Authors:** Gustavo Deco, Morten L. Kringelbach, Viktor K. Jirsa, Petra Ritter

## Abstract

In the human brain, spontaneous activity during resting state consists of rapid transitions between functional network states over time but the underlying mechanisms are not understood. We use connectome based computational brain network modeling to reveal fundamental principles of how the human brain generates large-scale activity observable by noninvasive neuroimaging. By including individual structural and functional neuroimaging data into brain network models we construct personalized brain models. With this novel approach, we reveal that the human brain during resting state operates at maximum metastability, i.e. in a state of maximum network switching. In addition, we investigate cortical heterogeneity across areas. Optimization of the spectral characteristics of each local brain region revealed the dynamical cortical core of the human brain, which is driving the activity of the rest of the whole brain. Personalized brain network modelling goes beyond correlational neuroimaging analysis and reveals non-trivial network mechanisms underlying non-invasive observations. Our novel findings significantly pertain to the important role of computational connectomics in understanding principles of brain function.

## INTRODUCTION

*“When we take a general view of the wonderful stream of our consciousness, what strikes us first is the different pace of its parts. Like a bird's life, it seems to be made of an alternation of flights and perchings.” William James ^1^*

Survival remains the perhaps most important problem faced by brains and a key challenge is how to segregate and integrate relevant information over different timescales when faced with hostile, often constantly changing environments ^2^. Reconciling different speeds of information processing, from fast to slow, is especially important, and could be key to the relative evolutionary success of mammals whose sophisticated brains are able to combine prior information from past memories with current stimuli to predict the future and to adapt behaviour accordingly ^3-5^.

This was recognized well over a century ago by William James, generally acknowledged as one of the fathers of modern cognitive psychology ^1^. Speaking of this problem using the apt metaphor of the stream of consciousness, James noted that there is a different pace to its parts, comparing it to the life of a bird whose journey consists of an “alternation of flights and perchings”. In the language of today's dynamical systems, the flights are akin to fast, segregative tendencies and the perchings to slower, integrative tendencies of the dynamic brain in action ^2,6,7^. In addition, motivated by recent experimental and modelling work of other labs ^8,9^, we investigate cortical heterogeneity across areas. By optimizing the spectral characteristics of each local brain node (in the coupled network), this allowed us to discover a dynamical core of the brain, i.e. the set of brain regions, which through their oscillations are driving the rest of the brain. Furthermore, with regards to balancing the different speeds of processing, a large body of psychological research has focused on what is known as dual process theories ^10,11^, identifying competing fast and slow systems which have to co-exist and function on multiple time-scales in order for the brain to efficiently allocate the resources necessary for survival ^12,13^.

Yet, the temporal dynamics and underlying neural mechanisms of this temporal processing on multiple timescales are poorly understood. Here we aim to provide a better understanding of the dynamics using computational brain network modelling which has emerged as a powerful tool for investigating the *causal* dynamics of the human brain, when carefully constrained by functional (FC) and structural connectivity (SC) obtained from empirical neuroimaging data ^14-18^. This theoretical framework has been largely successful in explaining the highly structured dynamics arising from spontaneous brain activity in the so-called resting-state-networks (RSN) ^19-21^, even if the resting brain never truly rests ^20^. Efficient task-related brain activity has been shown to rely on metastability of spontaneous brain activity allowing for optimal exploration of the dynamical repertoire ^22^ but it is not known if this metastability is maximally metastable ^6^. Our definition of metastability is taken from the work of Shanahan et al. as a measure of the variability of the Kuramoto order parameter (synchronization).

We investigated the dynamics of the brain network system through a local node neural mass description based on the most general form of expressing both noisy asynchronous dynamics and oscillations, namely a normal form of a Hopf bifurcation ^23-25^. Previous research has shown the usefulness, richness and generality of this type of model for describing EEG dynamics at the local node level ^24,25^. This normal form allowed us to fit the model to neuroimaging data *over time*, i.e. not only by fitting the grand average FC but also by fitting the temporal structure of the fluctuations, functional connectivity dynamics ^26^ (FCD, Figure 1A,B).

**Figure 1.**
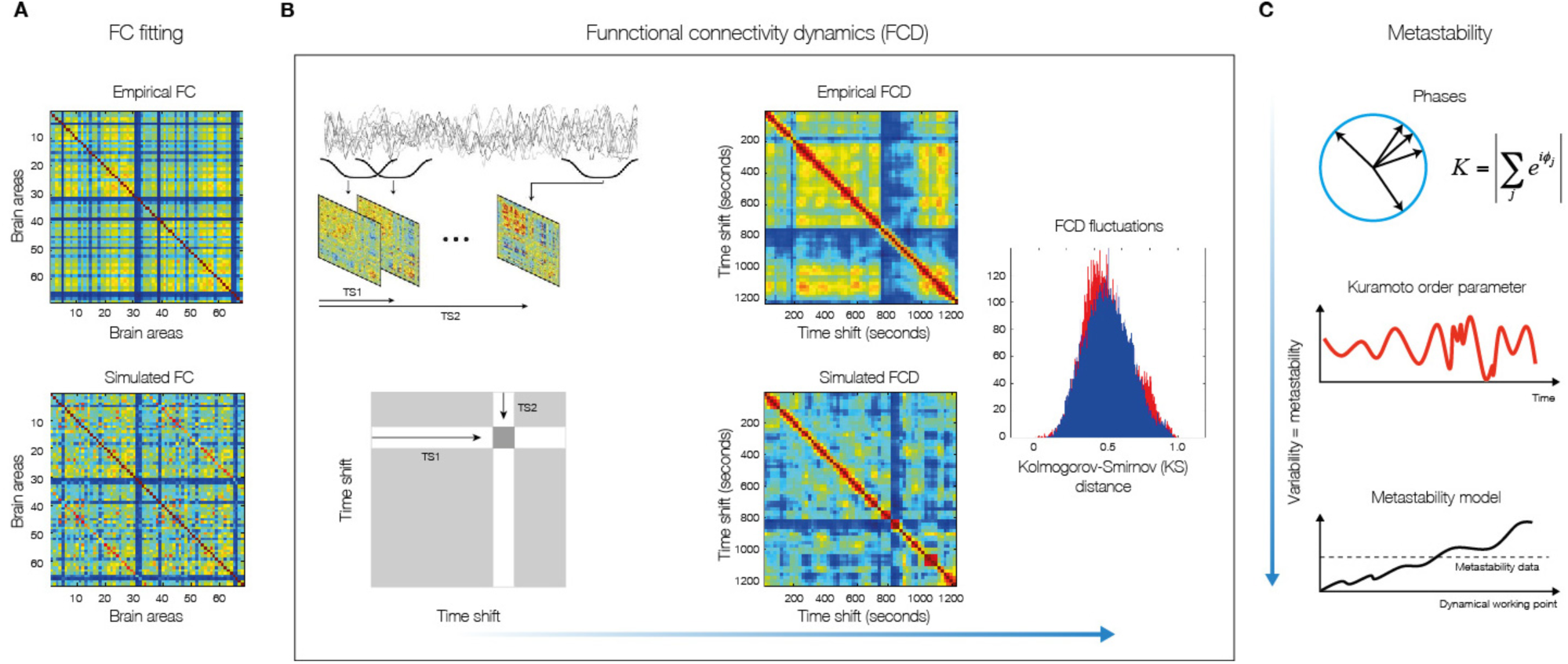
*Methods for measuring fit between simulated and empirical data. **A)** The fitting of the FC is measured by the Pearson correlation coefficient between corresponding elements of the upper triangular part of the matrices. **B)** For comparing the FCD statistics, we collected the upper triangular elements of the matrices (over all participants or sessions) and compared the simulated and empirical distribution by means of the Kolmogorov-Smirnov distance between them. The Kolmogorov–Smirnov distance quantifies the maximal difference between the cumulative distribution functions of the two samples. **C)** We measure the metastability as the standard deviation of the Kuramoto order parameter across time. The Kuramoto order parameter measures the global level of synchronization of the n oscillating signals. Under complete independence, the n phases are uniformly distributed and thus R is nearly zero, whereas R=1 if all phases are equal (full synchronization). For calculating the metastability of the empirical and simulated BOLD signals, we first band-pass filtered within the narrowband 0.04–0.07Hz and computed the instantaneous phase φ_k_(t) of each narrowband signal k using the Hilbert transform. The Hilbert transform yields the associated analytical signals. The analytic signal represents a narrowband signal, s(t), in the time domain as a rotating vector with an instantaneous phase, φ(t), and an instantaneous amplitude, A(t). Bottom panel visualizes a single example scenario (of many possible others) where the model system's metastability increases as a function of G. We also indicate the metastability measured in empirical data.*

We further explored if the optimal working point where FC and FCD are fitted corresponds to a dynamical region where the global metastability of the whole brain is maximized ^6^. In addition, motivated by recent experimental and modelling work of other labs ^8,9^, we investigate cortical heterogeneity across areas. By optimizing the spectral characteristics of each local brain node (in the coupled network), this allowed us discovering a dynamical core of the brain, i.e. the set of brain regions, which through their oscillations is driving the rest of the brain. As such this investigation was designed to provide an empirical, scientific footing for James’ metaphorical speculations of the flights and perchings of human brain dynamics, and to demonstrate the potential of sophisticated brain network computational modelling to provide new insights into the causal mechanisms of neuroimaging results.

## RESULTS

The results arose from using personalized brain network computational models for the analysis of empirical neuroimaging data characterising the functional and structural connectivity of 24 healthy human participants acquired using standard MRI techniques ^17^ (see *Methods*). In particular, we were able to gain new insights on the emergence of transiently spatiotemporal structured networks among segregated brain regions by examining a whole-brain network model using a very general neural mass model known as the normal form of a Hopf bifurcation (also known as Landau-Stuart Oscillators), which is the canonical model for studying the transition from noisy to oscillatory dynamics ^23^ (Figure 3). Here, we extended previous research on local node dynamics ^24,25^ by studying the whole-brain network dynamics, i.e. by investigating how those local noisy oscillators interact, and how the emerging whole-brain network activity relates to fMRI resting state dynamics. Within this model, each node of the network is modeled by a normal Hopf bifurcation, with an intrinsic frequency *ω_i_* in the 0.04–0.07Hz band (*i*=1, …,*n*). The intrinsic frequencies were estimated directly from the data, as given by the averaged peak frequency of the narrowband BOLD signals of each brain region (see *Methods*). The state of each node *i* is determined by its phase, *φ_i_*(*t*), and the interaction between nodes depends both on the structural couplings and the phase difference between the nodes. The model has only two types of control parameters, namely: one single global parameter, *G*, that represents the global scaling of the anatomical connectivity matrix, and the bifurcation parameters *a_j_* for each node (see Figure 3 and methods for the general structure and strategy of the brain network model).

**Figure 2.**
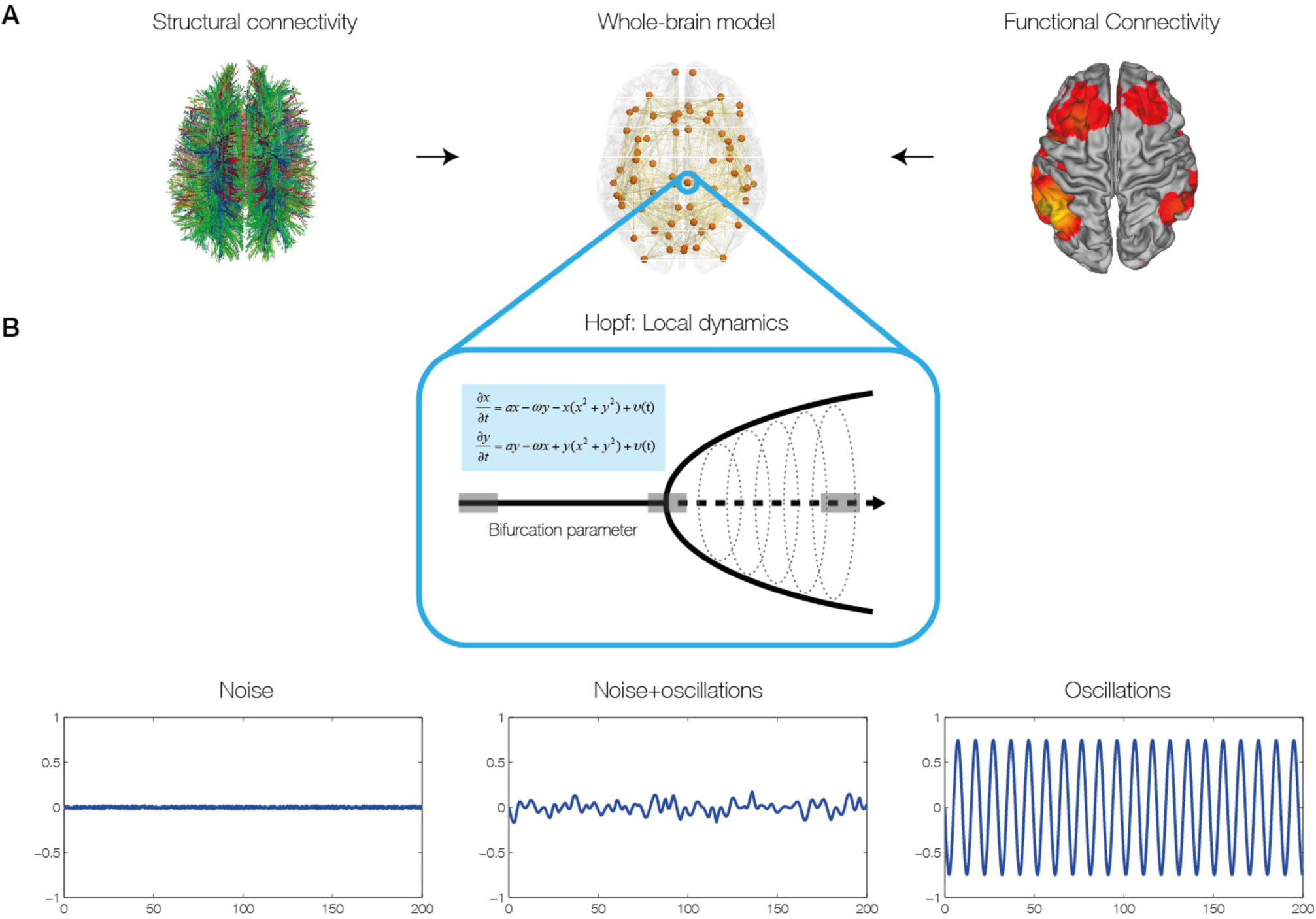
*Construction of individual brain network models. **A)** The brain network model was based on individual structural connectivity (SC) matrices from 24 participants derived from tractography of DTI (left) between the 68 regions of the Desikan-Kahilly parcellation (middle). The control parameters of the models were tuned using the grand average FC and FCD derived from fMRI BOLD data (right). **B)** For modelling local neural masses we used the normal form of a Hopf bifurcation, where depending on the bifurcation parameter, the local model generates a noisy signal (left), a mixed noisy and oscillatory signal (middle) or an oscillatory signal (right). It is at the border between noisy and oscillatory behaviour (middle), where the simulated signal looks like the empirical data, i.e. like noise with an oscillatory component around 0.05 Hz.*

### Maximal metastability at the optimal working point of model

Using the Hopf model, we were able discern the dynamical properties of the optimal working point of the system that is able to fit the characteristics of the empirical fMRI data. We were able to distinguish the origin of resting activity between the two hypothesized scenarios, namely: 1) noisy excursions at the edge of a critical bifurcation ^19,20,27,28^ or 2) metastable oscillations ^16^. The first scenario refers to the entrainment of noisy dynamics through the underlying anatomical connectivity matrix, i.e. inducing correlations of the local noise because of the underlying SC connections. The second scenario refers to the structuring of metastable cluster synchronizations of the underlying local oscillatory dynamics through the underlying anatomical SC connections. We define metastability as the standard deviation of synchrony at the network level described by order parameter R(t), where R(t) measures the phase uniformity and varies between 0 for a fully desynchronized network and 1 for a fully synchronized network (see methods and Figure 1C) ^29^. The present model is able to describe both types of dynamics, and the smooth transitions from one to the other, i.e. the transition from noisy to oscillatory dynamics (Figures 2). In order to distinguish the dynamical scenario, we investigated the capabilities of the model for fitting the grand average FC and also the time dependent characteristics of the RSN as reflected in the FCD in the different dynamical working regions (i.e. as a function of the control parameters). The grand average FC describes the mean spatial structure of the resting activity, whereas the FCD captures the statistical characteristic of the temporal structure of those spatial correlations (see *Methods* and ^26^).

Figure 3 shows that the best fit to the empirical data of Hopf model is found at the brink of the Hopf bifurcation. We equalized all local bifurcation parameters to a common value i.e. *a_j_* = *a*, in order to reduce the investigations to just two parameters, namely global bifurcation parameter *(a)* and global coupling strength *(G)*. Figure 3 shows how the empirical data are fitted in the Hopf model for different working points. The right column of Figure 3 shows the level of fitting of the FC, FCD and metastability. As can be seen, the best fitting of the three measures is obtained at the region on the brink of the Hopf bifurcation, i.e. for bifurcation parameter *a*, at the edge of zero on the negative side, such that the oscillators remain damped still. In this region not only the correlation between the empirical and simulated FC is maximized, but also the statistics of the rapid switching between FC(t) across time (FCD) is minimized in Kolmogorov-Smirnov sense, and the level of metastability of the data is reproduced. The fitting of the FC was measured by the Pearson correlation coefficient between corresponding elements of the upper triangular part of the matrices (see Figure 1 and *Methods*). For comparing the FCD statistics, we collected the upper triangular elements of the matrices (over all participants or sessions) and compared the simulated and empirical distribution by means of the Kolmogorov-Smirnov distance between them (see *Methods*).

**Figure 3.**
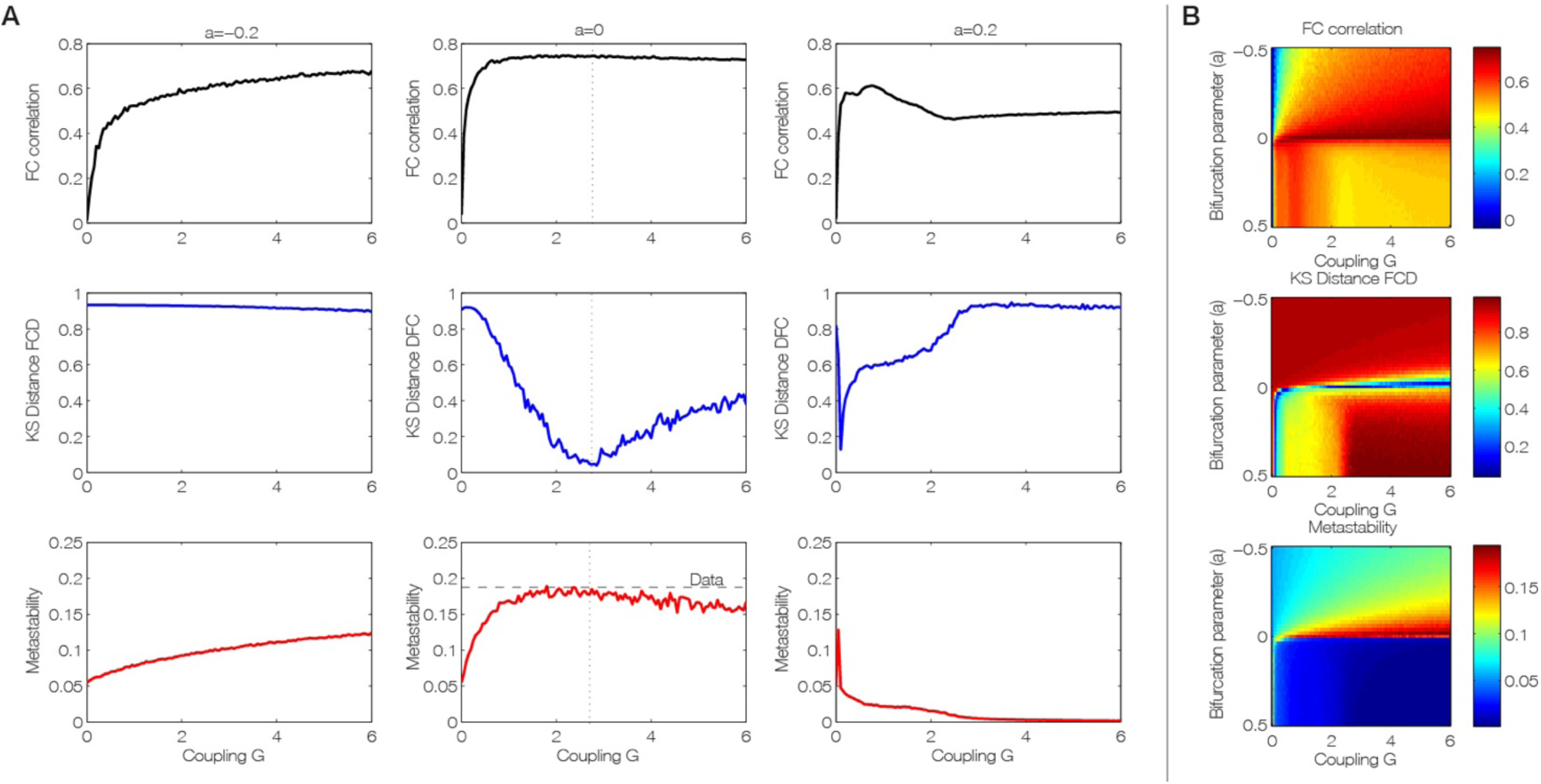
*Fitting of the empirical data by the brain network Hopf model for different working points. **A)** Level of fitting of the FC, FCD and metastability as a function of the global scaling parameter G for three different bifurcation parameters a=[−0.2 0 0.2], namely at the noisy oscillatory region, at the edge of the bifurcation and at the oscillatory regime. **B)** The three measures for assessing fitting between simulated and empirical data are shown color-coded as a function of bifurcation parameter a and global scaling parameter, G. The best fitting of the three measures is obtained for a region at the brink of the Hopf bifurcation, i.e. for bifurcation parameter a, at the edge of zero on the negative side. In this region not only the correlation between the empirical and simulated FC is maximized (upper panel), but also the statistics of the rapid switching between FC(t) across time (FCD) is minimized in Kolmogorov-Smirnov sense (middle panel), and the level of metastability of the data is perfectly reproduced (bottom panel*).

Furthermore, the results showed that only in the region at the border between noisy and oscillatory behaviour, is where the signals resembles the data, i.e. like noise with an oscillatory component around 0.05 Hz (Figure 2). The first three columns of Figure 3 show the dependence of those measurements as a function of the global scaling parameter *G* for three specific values of the bifurcation parameter *a*, namely at the noisy region, at the edge of the bifurcation and at the oscillatory regime. Clearly, the best results are obtained for the second column (at the edge of the bifurcation). The same panel shows that the FCD is the best constraining measure. There is a broad range of G where the FC and the metastability is well fitted, but only a relative narrow range where the FCD statistics is minimal, i.e. maximally fitted. In other words, the spatiotemporal structure of the FC is more informative than the grand average of the FC (i.e. the “classical” RSN). This is important, because until now, brain network models have always been fitted with the grand average FC - but see also ^26^.

We would like to remark that Figure 3 characterizes some of the bifurcation behaviour of the whole system. Indeed, the metastability for example serves as a network metric and characterizes the variability of this global synchronization as a function of those two control parameters. All three parameter spaces in Figure 3B, in conjunction, present a full picture of the spatiotemporal organization of the system. The three metrics characterize computationally the bifurcation properties of the full network dynamics.

Perhaps most importantly, as shown in Figure 3, the brain network model shows maximal *metastability* at the optimal working point of the model (*a*=0 and *G*=2.85), where the metastability is reflecting the variability of the synchronization between different nodes, i.e. the fluctuations of the states of phase configurations as a function of time ^29^. Further characterisation of these results is shown in Figure 4 which shows the optimal working point at the edge of the Hopf bifurcation (i.e. bifurcation parameter *a*=0), the FC, FCD and FCD statistics for three levels of global coupling *G* namely low, optimal and large. For comparison, the same matrices and distributions are plotted on the rightmost column for the empirical data (Figure 4B). Only the FCD and its statistics (bottom row) are constraining enough for optimizing the working point. Please note that for low *G* the FCD statistics does not show any switching between states in the RSN and that for very large *G* there are too much switching between states.

**Figure 4.**
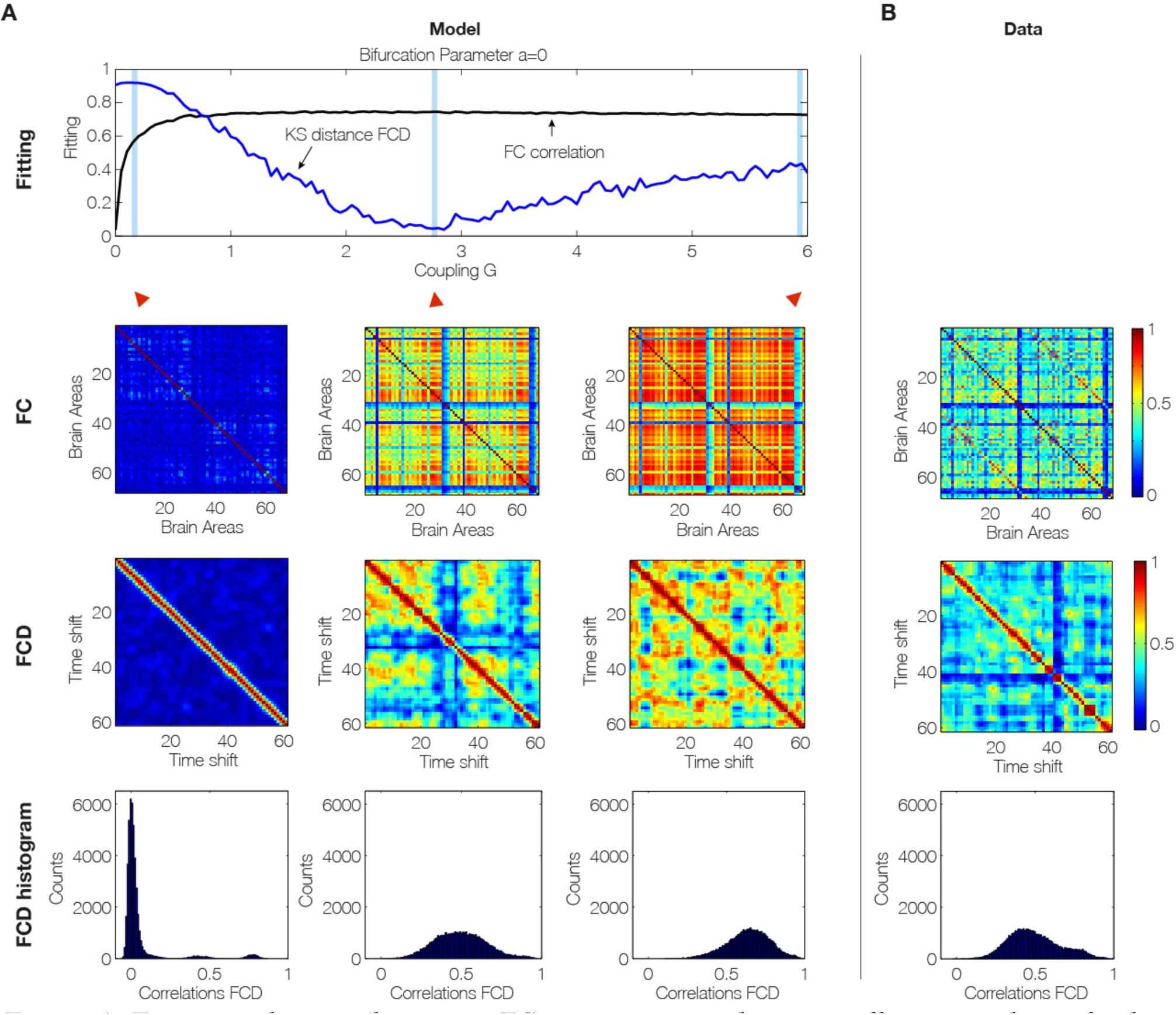
*Fitting to the grand average FC is a necessary but not sufficient condition for best empirical fitting. **A)** The figure shows the result of fitting the model to the empirical as a function of the global coupling parameter, G, at the optimal working point at the edge of the Hopf bifurcation (i.e. bifurcation parameter a=0). Three different coupling points were selected (low, optimal and large in the three columns) and we show the resulting FC correlation, FCD correlations and FCD histogram. Note that for low G the FCD statistics does not show any switching between RSN and that for very large G there are too much switching between states. **B)** For comparison, the same matrices and distributions are plotted for the empirical data. Note how only the FCD (row 2) and its statistics (row 3) are constraining enough for optimizing the working point the model to fit the empirical data (compare the distributions in row 3 and compare plots for FCD and FC fitting in row 1*).

### Dynamical core: contribution of individual brain regions to dynamics

In order to obtain information about the dynamical characteristics of each single brain area and to generate a heterogeneous brain network model (i.e. with different dynamics at each node), we optimized each single bifurcation parameter *a_j_* independently by fitting for each value of global coupling *G* the spectral characteristics of the simulated and empirical BOLD signals at each brain area (see *Methods*). The main results are plotted in Figure 5, where Figure 5a shows the evolution of the fitting of the FC and FCD statistics as a function of *G*. For large enough value of the global coupling a good fitting of both is obtained, i.e. large correlation between the empirical and simulated grand average FC and low difference in the statistics of the empirical and simulated FCD (Kolmogorov-Smirnov distance). Please note that Figure 5a is generated in a different way than Figure 4A (which uses only the optimum fit a=0 for all regions and G=2.85). Instead, in Figure 5A, for each G we optimize the bifurcation value, *a*, for *each* region (shown in Figure 5b). As can be seen at a critical value of G, the bifurcation values remain the same, only scaled. Thus the FCD fit in Figure 5A will asymptote as G increases.

**Figure 5.**
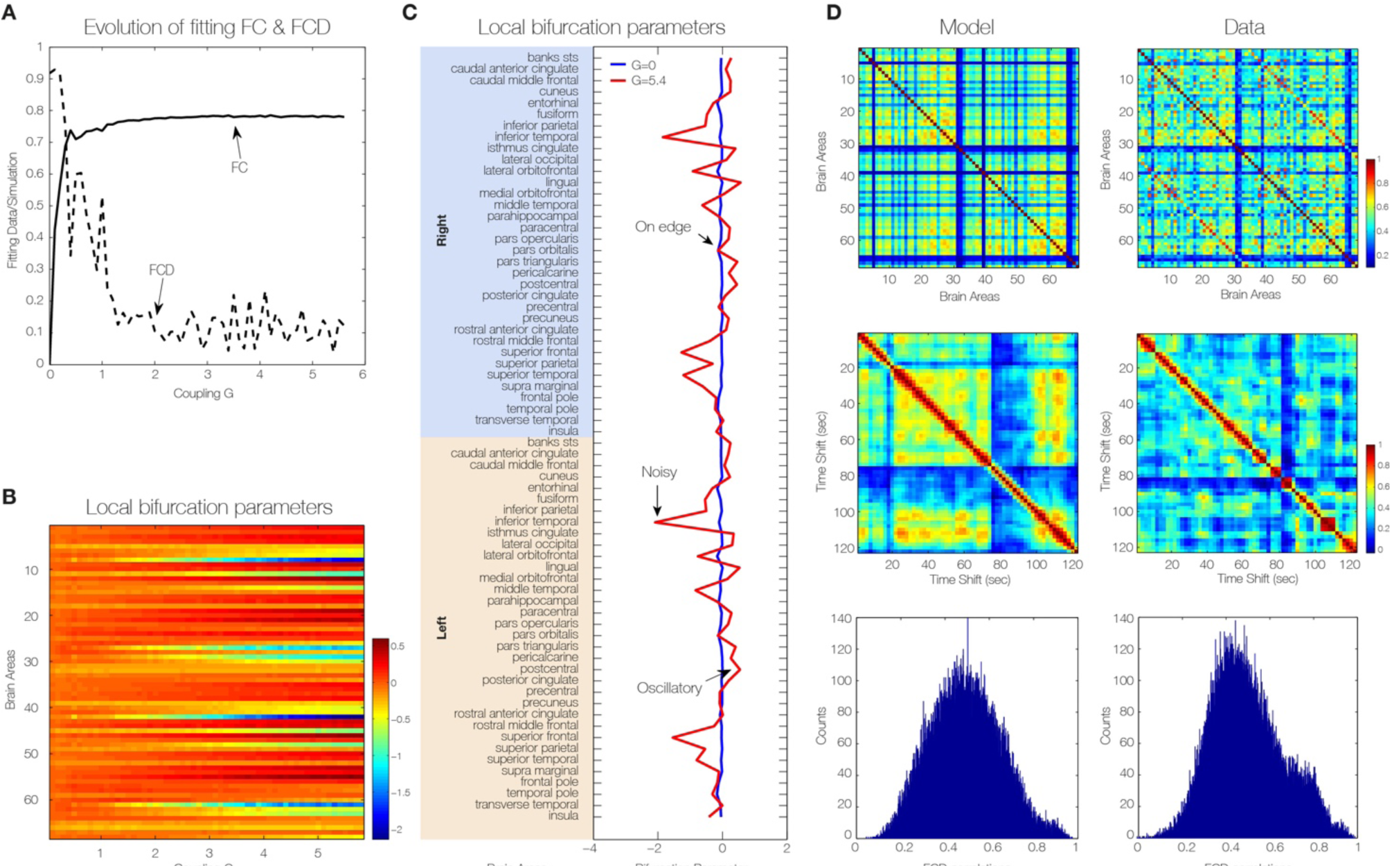
*Spectral characteristics of the dynamical core of the human brain. To generate a heterogeneous brain network model (i.e. with different dynamics at each node), we optimized each single bifurcation parameter a_j_ independently by fitting for each value of global coupling G the spectral characteristics of the simulated and empirical BOLD signals at each brain area. **A)** The evolution of the fitting of the FC and FCD statistics as a function of G. For large enough value of the global coupling a good fitting of both is obtained, i.e. large correlation between the empirical and simulated grand average FC and low difference in the statistics of the empirical and simulated FCD (Kolmogorov-Smirnov distance). **B)** The evolution of the single values of the local bifurcations parameters a_j_ as a function of the global coupling G. For low values of G homogeneous local bifurcation parameters a_j_ around zero are obtained. When the level of fitting improves for larger values of G a more heterogeneous distribution of a_j_ is obtained. **C)** The local bifurcation parameters for each region for the uncoupled network (i.e. G=0) and for the optimal coupling (G=5.4). If the network is uncoupled, each single brain area fitted the spectral characteristics of the empirical BOLD signals in a very homogeneous way by local bifurcations parameters at the edge of the local Hopf bifurcation, i.e. at zero. **D)** When the whole-brain network is coupled, we can discover the “true” intrinsic local dynamics that fits the local empirical BOLD characteristics and the global quantities FC, FCD and metastability.*

For optimizing *a_j_* values, we use a greedy optimisation strategy, where we iteratively increase or decrease the *a_j_* value according to the local power of the signal in a given region *j*. Greedy algorithms exploit local optima, but often approximate optimal solutions well in reasonable time and produce good results as shown in Figure 5. The local bifurcation parameters for each region for the uncoupled network (i.e. *G*=0) and for the optimal coupling (*G*=5.4) can be seen in Figure 5c. If the network is uncoupled, each single brain area fitted the spectral characteristics of the empirical BOLD signals in a very homogeneous way by local bifurcations parameters at the edge of the local Hopf bifurcation, i.e. at zero. When the brain network is coupled, the “true” intrinsic local dynamics for the profile of optimal local bifurcation parameters *a_j_* observed at that point that fit the local empirical BOLD characteristics and the global quantities FC, FCD and metastability (Figure 5d).

Brain regions, for which best predictions were achieved in an oscillatory mode, i.e. with bifurcation parameters *a* > 0.1 are visualised in Figure 6. We found that the dynamical core within this parcellation consisted of eight lateralised brain regions: medial orbitofrontal cortex, posterior cingulate cortex and transverse temporal gyrus in the right hemisphere, and caudal middle frontal gyrus, precentral gyrus, precuneus cortex, rostral anterior cingulate cortex and transverse temporal gyrus in the left hemisphere. Those nodes working at the edge of the bifurcation are highlighted as a "dynamical core" whose perturbations can propagate in an optimal way to the rest of the network.

**Figure 6.**
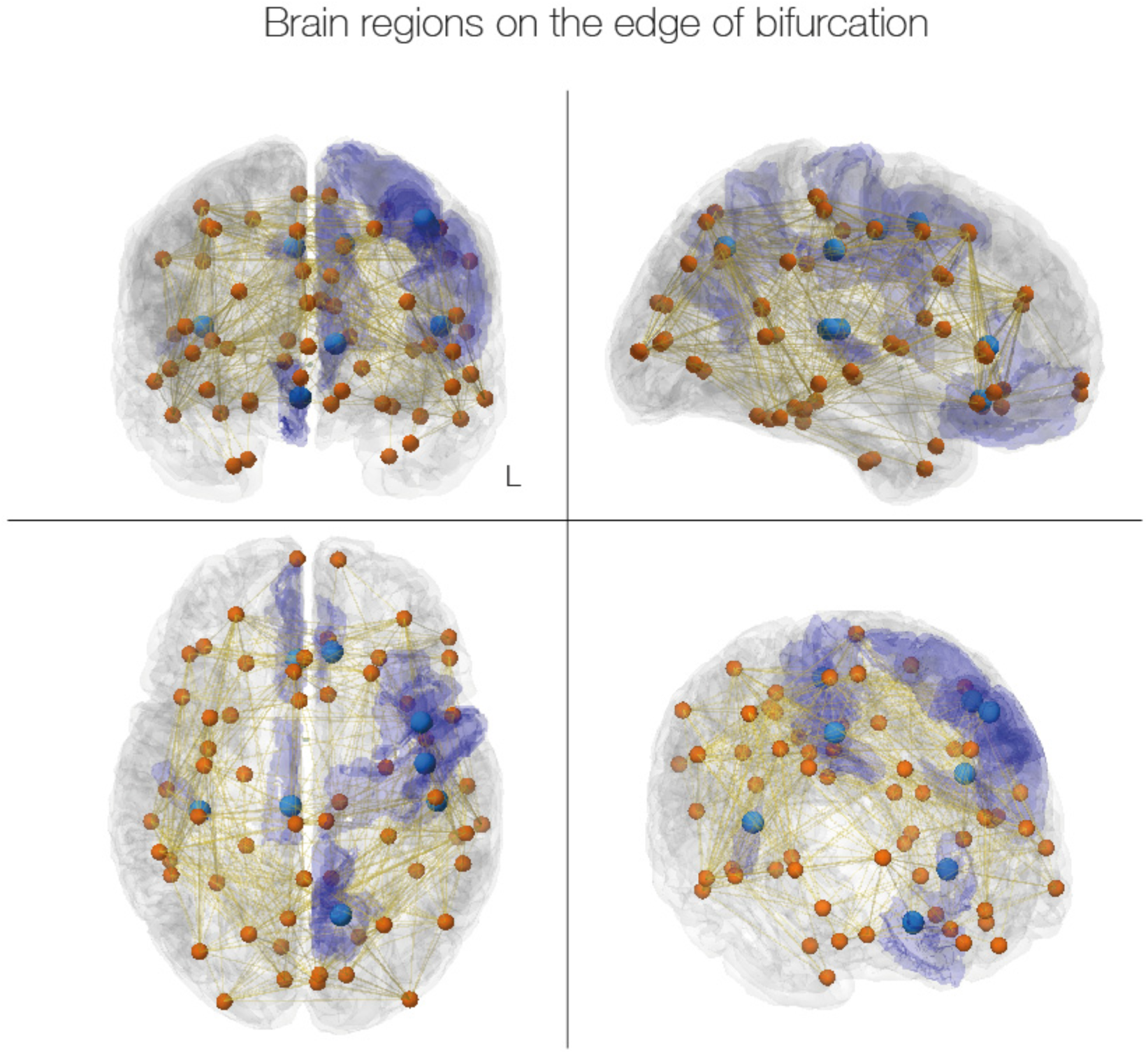
*Dynamical core in the human brain. The figure shows the dynamical core regions on the edge of bifurcation (location of neural masses shown in light blue and transparent blue for the full region). These are the nodes with the ability to react immediately to changes in the predicted input and thus likely to drive the rest of the brain networks. The eight regions are clearly lateralised; and in the right hemisphere encompass medial orbitofrontal cortex, posterior cingulate cortex and transverse temporal gyrus, while in the left hemisphere include caudal middle frontal gyrus, precentral gyrus, precuneus cortex, rostral anterior cingulate cortex and transverse temporal gyrus. Interestingly, some of these regions are part of the default mode network (medial orbitofrontal cortex, posterior cingulate cortex and precuneus cortex) while others have been implicated in memory processing (parahippocampal and transverse temporal gyrus), auditory processing (transverse temporal gyrus), selection for action (rostral anterior cingulate cortex and caudal middle frontal gyrus) and motor execution (precentral gyrus*).

## DISCUSSION

We provide mechanistic explanations of the complex spatiotemporal dynamics of brain function arising from James’ early speculations ^1^ to much more detailed scientific enquiry ^2,30-32^. This confirms that brain results from complex interactions in a system of non-linearly coupled, non-linear oscillatory processes which display dynamical system phenomena such as multiple stable states, instability, state transitions and metastability, of which the latter has been proposed to form a core dynamical description of coordinated brain and behavioral activity ^6^.

In the 1980s the physicist Hermann Haken suggested to mechanistically interpret brain processes of segregation and integration as a sequence of semistable states, so-called saddle states ^33^. He proposed to view the complex integrative and segregative tendencies as expressions of emergent lower-dimensional behavior of collective variables, which he termed ‘order parameters’. Scott Kelso popularized this concept using the term ‘metastability’ based on his brain-behaviour experiments and drawing inspiration from other researchers including Rodolfo Llinás and Francisco Varela ^31,32^. He generalized metastability to include the oscillatory states of brain processes found in between complete synchronization and independence ^30,34^. Later research has formalized these concepts more rigorously, e.g. via the heteroclinic channel 35,36 and Structured Flows on Manifolds (SFM) ^37,38^.

We shed new causal light on the mechanisms underlying RSNs by extending previous research which has demonstrated the existence of RSNs, i.e. brain networks correlated within the grand average FC during resting state ^21,39,40^. FC has become routinely used as a biomarker in various clinical applications, even though its predictive value holds only for group analyses, and not currently for the individual ^41^. This problem arises most likely from the lack of taking time into account, i.e. the non-stationary nature of the resting state dynamics ^42,43^. Hansen and colleagues demonstrated that the grand average FC is more closely linked to the SC and linear models of FC ^26^. When non-linearities are considered in the network models, the spatiotemporally dynamic repertoire of the network is significantly enhanced and the resting state dynamics shows the non-stationary FCD, which expresses itself as the switching dynamics of the FC. While Hansen and colleagues proposed FCD as a novel biomarker and demonstrated that all known RSNs can be derived from the non-linear network dynamics of FCD, they did not fit the model to the empirical functional time series data. The patterns in the FCD matrix arise from what is essentially a random process and thus different for different measurements. This renders the fitting process for brain network models more complex than fitting with the grand average FC, for which a Pearson correlation across empirical and simulated FC matrices is sufficient.

We have addressed this issue through a systematic fitting approach of the random process in FCD to the empirical data. The conjunction of using sophisticated fitting and systematic parameter analysis allowed us to test the mechanistic hypotheses underlying the resting state, i.e. whether the brain at rest operates close to the edge of a bifurcation and/or occupies a metastable state. Both scenarios can be mechanistically realized by non-linearly coupling Hopf bifurcators ^23^. Hopf oscillators have been used previously in connectome-based modelling of resting state dynamics in EEG/MEG and fMRI ^14^, as well as for the modelling of the detailed temporal dynamics in EEG/MEG ^24,25^. The usage here though is different from the previous research, since the Hopf oscillators act as the sources of BOLD signal in the connectome based network model. Ghosh and colleagues used the Hopf oscillators as the sources of the electrophysiological signal and employed the Balloon Windkessel to derive the BOLD signal ^44^. Given this interpretation, they needed to include all the signal transmission delays. In our present approach, the oscillation frequencies are significantly slower and thus permit the neglect of the time delays, which simplifies the computational effort of the simulation and thus the computational fitting of the models against empirical data.

Our key finding is the demonstration that the optimal operating regime is at the edge of the local Hopf bifurcation, i.e. a balance of noisy excursions in the oscillatory state. We not only were able to demonstrate that previous findings on the optimal operating point based on grand-average FC hold true if we take into account the temporal dynamics of FC, i.e. FCD. We also demonstrated that a better way of constraining brain network models is by not only fitting the grand average FC, but by also fitting the temporal structure of the fluctuations using the FCD.

Another remarkable and important finding is that high metastability is only present in a narrow range of bifurcation parameter when *a* is close to the edge of the bifurcation. In other words, the FCD of the spontaneous resting state, in conjunction with brain network modelling provide evidence that the brain at rest is maximally metastable, refining and demonstrating the hypothesis of Tognoli and Kelso ^6^. Note that there is also a region for very small *G* and positive *a* (oscillatory regime) where a relatively good fitting is obtained. This dynamic regime was previously observed with a pure oscillatory Kuramoto model of the BOLD signals at the mesoscopic level ^45^. Nevertheless, the level of fitting for the FC, metastability and even FCD is not as good as the one obtained in the region at the edge of the Hopf bifurcation. On the other hand, besides the extreme sensitivity of that working point (ultra-narrow regime of optimality) which means that the result is not so robust, the qualitative description of the BOLD signals is not realistic in the pure oscillatory regime in comparison with the noisy/oscillatory excursions evidenced in the regime of the bifurcation parameter *a* near zero.

For constructing a heterogeneous brain network model with different local parameter values, we took into account the spectral information of the BOLD data. We addressed the question if the oscillations at the individual nodes play a mechanistic role for the emergence of FC/FCD. In particular, we identified a cortical core of eight brain regions with the optimal fit of bifurcation parameter *a* close to the edge of bifurcation. We propose to call this the *dynamical cortical core* of the brain. Interestingly, three of these regions (the medial orbitofrontal cortex, posterior cingulate cortex and precuneus cortex) are part of the default mode network and thus re-experience past events and pre-experience possible future events ^46,47^. In this vein other regions (parahippocampal and transverse temporal gyrus) have also been implicated in memory processing and may thus perhaps be helping integrate information over different timescales, binding fast and slow processes over time ^2^. This information is always contextual and in the noisy, unpredictable scanner it is perhaps not surprising that the brain is attending to the auditory signals (transverse temporal gyrus). As such this information processing is available for conflict monitoring and selection for action (rostral anterior cingulate cortex and caudal middle frontal gyrus) and motor execution (precentral gyrus) ^48^. Equally, the involvement of the cingulate cortex is interesting given that this region recently has been shown to be part of the common neurobiological substrate for mental illness across across six diverse diagnostic groups (schizophrenia, bipolar disorder, depression, addiction, obsessive-compulsive disorder, and anxiety) based on a meta-analysis of grey matter loss in 193 neuroimaging studies of 15892 individuals ^49^. This reinforces the potential use of brain network computational modelling for understanding the underlying mechanisms of neuropsychiatric disorders ^50^. The right-handed quality of Figure 6, presumably arises from the specifics of the data used to fit the model and we will be exploring its biological validity in subsequent studies with larger group sizes.

Although the bifurcation parameter does not have a direct biophysical correlate, it seems to be involved in mediating biophysical effects. We therefore propose that in future both the global bifurcation parameter as well as the individual parameters could potentially serve as biomarkers for disease. It will be important to explore the changes for different brain diseases, e.g. within a standardized framework for connectome-based modelling such as The Virtual Brain (TVB) ^51,52^, and applications such as fitting of TVB's dynamic regime and TVB Processing pipeline ^17^.

Overall, we have shown that neuroimaging data can be causally analysed by constructing a relatively simple brain network computational model using a Hopf bifurcation. This model was shown to be maximally metastable at the optimal fitting with the spatiotemporal dynamics of spontaneous brain activity. This dynamical regime may well allow for the optimal integration and segregation of fast and slow information over different time-scales, the “flights” and “perchings” of the stream of consciousness alluded to by William James over 100 years ago.

## METHODS

### Ethics Statement

All participants of this study gave written informed consent before the study, which was performed in compliance with the relevant laws and institutional guidelines and approved by the ethics committee of the Charité University Berlin.

### Empirical MRI Data Collection

Structural data from DTI and resting-state BOLD signal time series were acquired for 24 healthy participants (age between 18 and 33 years old, mean 25.7, 12 females, 12 males). A full description of the generation of SC and FC matrices from those data can be found in ^17^. Here, we provide a quick overview of the employed methods. Empirical data were acquired at Berlin Center for Advanced Imaging, Charité University Medicine, Berlin, Germany. For simultaneous EEG-fMRI ^53,54^, participants were asked to stay awake and keep their eyes closed. No other controlled task had to be performed. In addition, a localizer, DTI and T2 sequence were recorded for each participant. MRI was performed using a 3 Tesla Siemens Trim Trio MR scanner and a 12-channel Siemens head coil. Specifications for the employed sequences can be found in ^54^. For each participant anatomical T1-weighted scans were acquired. DTI and GRE field mapping were measured directly after the anatomical scans. Next, functional MRI (BOLD-sensitive, T2*-weighted, TR 1940 ms, TE 30 ms, FA 78°, 32 transversal slices (3 mm), voxel size 3 x 3 x 3 mm, FoV 192 mm, 64 matrix) was recorded simultaneously to the EEG recording.

### MRI Data Analysis

Processing steps executed by the public Berlin automatized processing pipeline ^54^ comprised 1) preprocessing of T1-weighted scans, cortical reconstruction, tessellation and parcellation, 2) transformation of anatomical masks to diffusion space, 3) processing of diffusion data, 4) transformation of anatomical masks to fMRI space, 5) Processing of fMRI data

### Anatomical MRI Data Analysis

The highly resolved anatomical images are important to create a precise parcellation of the brain. For each of those parcellated units, empirical functional data time series are spatially aggregated. T1-weighted images are pre-processed using FREESURFER including probabilistic atlas based cortical parcellation, here using Desikan-Killany (DK) atlas ^55^ (Table 1). This generates volumes that contain all cortical and subcortical parcellated regions with corresponding region labels used for fiber-tracking and BOLD time-series extraction.

**Table 1.** *Anatomical labels for the 68 regions in the Desikan-Kahilly parcellation.* The two region numbers per line refer to right and left hemisphere respectively.

| **Region number** | **Region name** |
| --- | --- |
| 1;35 | Superior temporal sulcus, banks of |
| 2;36 | Caudal anterior cingulate cortex |
| 3;37 | Caudal middle frontal gyrus |
| 4;38 | Cuneus cortex |
| 5;39 | Entorhinal cortex |
| 6;40 | Fusiform gyrus |
| 7;41 | Inferior parietal cortex |
| 8;42 | Inferior temporal gyrus |
| 9;43 | Isthmus of cingulate cortex |
| 10;44 | Lateral occipital cortex |
| 11;45 | Lateral orbitofrontal cortex |
| 12;46 | Lingual gyrus |
| 13;47 | Medial orbitofrontal cortex |
| 14;48 | Middle temporal gyrus |
| 15;49 | Parahippocampal gyrus |
| 16;50 | Paracentral lobule |
| 17;51 | Pars opercularis |
| 18;52 | Pars orbitalis |
| 19;53 | Pars triangularis |
| 20;54 | Pericalcarine cortex |
| 21;55 | Postcentral gyrus |
| 22;56 | Posterior cingulate cortex |
| 23;57 | Precentral gyrus |
| 24;58 | Precuneus cortex |
| 25;59 | Rostral anterior cingulate cortex |
| 26;60 | Rostral middle frontal gyrus |
| 27;61 | Superior frontal cortex |
| 28;62 | Superior parietal cortex |
| 29;63 | Superior temporal gyrus |
| 30;64 | Supramarginal gyrus |
| 31;65 | Frontal pole |
| 32;66 | Temporal pole |
| 33;67 | Transverse temporal cortex (primary auditory cortex) |
| 34;68 | Insula |

### Empirical DTI Data Analysis and Tractography

Tractography requires binary WM masks to restrict tracking to WM voxels. Upon extraction of gradient vectors and values (known as b-table) using MRTrix, dw-MRI data are pre-processed using FREESURFER. Besides motion correction and eddy current correction (ECC) the b0 image is linearly registered (6 degrees of freedom, DOF) to the participant's anatomical T1-weighted image and the resulting registration rule is stored for later use. We transformed the high-resolution mask volumes from the anatomical space to the participant's diffusion space, to further use it for fiber tracking. The cortical and subcortical parcellations are resampled into diffusion space, one time using the original 1 mm isotropic voxel size (for subvoxel seeding) and one time matching that of our dw-MRI data, i.e., 2.3 mm isotropic voxel size. During MRTrix pre-processing diffusion tensor images that store the diffusion tensor (i.e., the diffusion ellipsoid) for each voxel location are computed. Based on that, a fractional anisotropy (FA) and an eigenvector map are computed and masked by the binary WM mask created previously. For subsequent fiber-response function estimation, a mask containing high-anisotropy voxels is computed. Fibre orientation distributions are estimated using constrained spherical deconvolution ^56^ based on a response function estimated in voxels that are expected to contain a single, coherently-oriented fibre bundle (commands dwi2response tournier and dwi2fod; see MRTrix Documentation: http://mrtrix.readthedocs.io/en/latest/). In order to resolve crossing pathways, fibers are prolonged by employing a probabilistic tracking approach as provided by MRTrix. In order to exclude spurious tracks, three types of masks are used to constrain tracking: seeding-, target- and stop-masks. In order to restrict track-prolongation to WM, a WM-mask that contains the union of GM-WM-interface and cortical WM voxels is defined as a global stop mask for tracking. To address several confounds in the estimation of connection strengths (information transmission capacities), a new seeding and fiber aggregation strategy was employed developed for this pipeline and described in detail in ^17^. In combination with a new aggregation scheme, it is based on an appropriate selection of seed voxels and controlling for the number of generated tracks in each seed voxel. Instead of using every WM voxel, tracks are initiated from GM-WM-interface voxels and a fixed number of tracks are generated for each seed-voxel. Since a GM parcellation-based aggregation is performed, each seed-mask is associated with a ROI of the GM atlas. Along with seeding-masks complementary target-masks are defined specifying valid terminal regions for each track that was initiated in a specific seed voxel. The capacity measures that we derive between each pair of regions are intended to estimate the strength of the influence that one region exerts over another, i.e., their SC. In order to improve existing methods for capacities estimation the approach makes use of several assumptions with regard to seed-ROI selection, tracking and aggregation of generated tracks^17^. Upon tractography the pipeline aggregates generated tracks to structural connectome matrices. The weighted distinct connection count used here divides each distinct connection by the number of distinct connections leaving the seed-voxel (yielding asymmetric capacities matrix). Values have been normalized by the total surface area of the GWI of a participant.

### Empirical fMRI Data Analysis

In order to generate the FC matrices, FSL's FEAT pipeline is used to perform the following operations: deleting the first five images of the series to exclude possible saturation effects in the images, high-pass temporal filtering (100 seconds high-pass filter), motion correction, brain extraction and a 6 DOF linear registration to the MNI space. Functional data is registered to the participant's T1-weighted images and parcellated according to FREESURFER's cortical segmentation. By inverting the mapping rule found by registration, anatomical segmentations are mapped onto the functional space. Finally, average BOLD signal time series for each region are generated by computing the mean over all voxel time-series for each region. From the region wise aggregated BOLD data, FC matrices are computed within MATLAB using and Pearson's linear correlation coefficient as FC metrics. We did not perform global signal regression on data.

### Brain Network Model

The brain network model consists of 68 coupled brain areas (nodes) derived from the parcellation explained above. The global dynamics of the brain network model used here results from the mutual interactions of local node dynamics coupled through the underlying empirical anatomical structural connectivity matrix *C_ij_* (see Figure 2). The structural matrix *C_ij_* denotes the density of fibres between cortical area i and j as extracted from the DTI based tractography (scaled to a maximum value of 0.2). The local dynamics of each individual node is described by the normal form of a supercritical Hopf bifurcation, which is able to describe the transition from asynchronous noisy behavior to full oscillations. Thus, in complex coordinates, each node *j* is described by following equation:

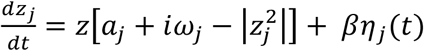

where

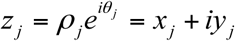

and *η_i_* (*t*) is additive Gaussian noise with standard deviation β=0.02. This normal form has a supercritical bifurcation at *a_j_* = 0, so that for *a_j_* < 0 the local dynamics has a stable fixed point at *z_j_* = 0 (which because of the additive noise corresponds to a low activity asynchronous state) and for *a_j_* > 0 there exists a stable limit cycle oscillation with frequency 
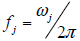
. We insert equation 2 in equation 1 and separate real part in equation 3 and imaginary part in equation 4. Thus, the whole-brain dynamics is defined by following set of coupled equations:

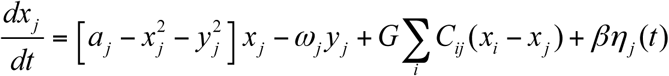

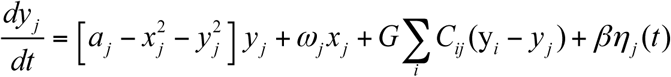

Please note that following the literature from physics, the equations are written in Cartesian, rather than polar coordinates ^57-60^, We couple the equations using the common difference coupling, which approximates the simplest (linear) part of a general coupling function. These equations are valid in the weakly coupled oscillator limit, in which the coupling preserves the periodic orbit of the uncoupled oscillators. If the linear coupling (following a Taylor expansion of the full coupling) does not exist, the next non-vanishing higher order term should be considered, which is a case we do not address here (please see Kuramoto ^57^ (see Eq 5.3.1) and Pikovsky, Arkady and Kurths ^58^ (see Eq. 8.12) for more detailed analytic treatments of the equations).

In the latter equations, G is a global scaling factor (global conductivity parameter scaling equally all synaptic connections). The global scaling factor G and the bifurcation parameters *a_j_* are the control parameters with which we study the optimal dynamical working region where the simulations maximally fit the empirical FC and the FCD. We model with the variables *x_j_* the BOLD signal of each node *j*. The empirical BOLD signals were band-pass filtered within the narrowband 0.04–0.07 Hz. This frequency band has been mapped to the gray matter and it has been shown to be more reliable and functionally relevant than other frequency bands ^61-64^. Within this model, the intrinsic frequency *ω_j_* of each node is in the 0.04–0.07Hz band (*i*=1, …,*n*). The intrinsic frequencies were estimated from the data, as given by the averaged peak frequency of the narrowband BOLD signals of each brain region.

### Grand average FC and FCD matrices

The grand average FC is defined as the matrix of correlations of the BOLD signals between two brain areas over the whole time window of acquisition. In order to characterize the time dependent structure of the resting fluctuations, we estimate the FCD matrix ^26^ (see Figure 1). Each full-length BOLD signal of 22 min is split up into M=61 sliding windows of 60 sec, overlapping by 40 sec. For each sliding window, centered at time t, we calculated a separate FC matrix, FC(t). The FCD is a MxM symmetric matrix whose (t1, t2) entry is defined by the Pearson correlation between the upper triangular parts of the two matrices FC(t1) and FC(t2). Epochs of stable FC(t) configurations are reflected around the FCD diagonal in blocks of elevated inter-FC(t) correlations.

The grand average FC and the FCD matrices were estimated for the recordings of each of the 24 participants as well as for 24 simulations of 22 minutes of the computational model. We compared the FC matrices of the model (averaged Fisher's z-transformed over the 24 sessions) and the empirical data (averaged Fisher's z-transformed over the 24 participants), adopting as a measure of similarity between the two matrices the Pearson correlation coefficient between corresponding elements of the upper triangular part of the matrices. For comparing the FCD statistics, we collected the upper triangular elements of the matrices (over all participants or sessions) and generated the distribution of them. Then, we compared the simulated and empirical distribution by means of the Kolmogorov-Smirnov distance between them. The Kolmogorov–Smirnov distance quantifies the maximal difference between the cumulative distribution functions of the two samples.

### Metastability

Here, we refer to metastability as a measure of how variable are the states of phase configurations as a function of time, i.e. how the synchronization between the different nodes fluctuates across time ^29^. Thus, we measure the metastability as the standard deviation of the Kuramoto order parameter across time. The Kuramoto order parameter is defined by following equation:

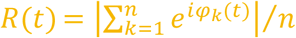

where φ_*k*_(*t*) is the instantaneous phase of each narrowband BOLD signal at node *k.* The Kuramoto order parameter measures the global level of synchronization of the *n* oscillating signals. Under complete independence, the *n* phases are uniformly distributed and thus *R* is nearly zero, whereas *R*=1 if all phases are equal (full synchronization). Thus, for calculating the metastability of the empirical and simulated BOLD signals, we first band-pass filtered within the narrowband 0.04– 0.07Hz (as previously explained) and computed the instantaneous phase φ_*k*_(*t*) of each narrowband signal *k* using the Hilbert transform. The Hilbert transform yields the associated analytical signals. The analytic signal represents a narrowband signal, *s*(*t*), in the time domain as a rotating vector with an instantaneous phase, *φ*(*t*), and an instantaneous amplitude, *A*(*t*), i.e., *s*(*t*) = *A*(*t*)cos *φ*(*t*). The phase and the amplitude are given by the argument and the modulus, respectively, of the complex signal *z*(*t*), given by *z*(*t*) = *s*(*t*) + *i*.H[*s*(*t*)], where *i* is the imaginary unit and H[*s*(*t*)] is the Hilbert transform of *s*(t).

### Local Optimization of Brain Nodes

The local optimization of each single bifurcation parameter *a_j_* is based on the fitting of the spectral information of the empirical BOLD signals in each node. In particular, we aim to fit the proportion of power in the 0.04-0.07 Hz band with respect to the 0.04-0.25 Hz band (i.e. we remove the smallest frequencies below 0.04 Hz and consider the whole spectra until the Nyquist frequency which is 0.25 Hz) ^45^. For this, we filtered the BOLD signals in the 0.04-0.25 Hz band, and calculated the power spectrum *P_j_* (*f*) for each node *j*. We define the proportion,

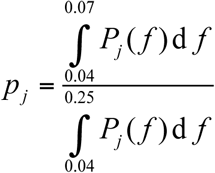

and update the local bifurcation parameters by a gradient descendent strategy, i.e.:

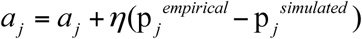

until convergence. We used here *η* = 0.1. The updates of the *a_j_* values are done in each optimization step in parallel.

## References

1 James, W. The Principles of Psychology. (Henry Holt, 1890).

2 Deco, G., Tononi, G., Boly, M. & Kringelbach, M. L. Rethinking segregation and integration: contributions of whole-brain modelling Nat Rev Neurosci 16, 430-439 (2015).

3 Berridge, K. C. & Kringelbach, M. L. Pleasure systems in the brain. Neuron 86, 646-664 (2015).

4 Friston, K. & Kiebel, S. Predictive coding under the free-energy principle. Philosophical transactions of the Royal Society of London. Series B, Biological sciences 364, 1211-1221, doi:10.1098/rstb.2008.0300 (2009).

5 Kringelbach, M. L., McIntosh, A. R., Ritter, P., Jirsa, V. K. & Deco, G. The rediscovery of slowness: exploring the timing of cognition. TICS 19, 616-628 (2015).

6 Tognoli, E. & Kelso, J. A. The metastable brain. Neuron 81, 35-48, doi:10.1016/j.neuron.2013.12.022 (2014).

7 Friston, K. J. The labile brain. I. Neuronal transients and nonlinear coupling. Philosophical transactions of the Royal Society of London. Series B, Biological sciences 355, 215-236, doi:10.1098/rstb.2000.0560 (2000).

8 Chaudhuri, R., Knoblauch, K., Gariel, M. A., Kennedy, H. & Wang, X. J. A Large-Scale Circuit Mechanism for Hierarchical Dynamical Processing in the Primate Cortex. Neuron 88, 419-431, doi:10.1016/j.neuron.2015.09.008 (2015).

9 Simony, E. et al. Dynamic reconfiguration of the default mode network during narrative comprehension. Nat Commun 7, 12141, doi:10.1038/ncomms12141 (2016).

10 Stanovich, K. E. & West, R. F. Individual differences in reasoning: implications for the rationality debate? The Behavioral and brain sciences 23, 645-665; discussion 665-726 (2000).

11 Posner, M. I. & Snyder, C. R. R. in Information processing and cognition: The Loyola Symposium (ed R. L. Solso) 55-85 (Wiley, 1975).

12 Tversky, A. & Kahneman, D. Judgment under Uncertainty: Heuristics and Biases. Science 185, 1124-1131, doi:185/4157/1124 [pii] 10.1126/science.185.4157.1124 (1974).

13 Kahneman, D. Thinking, fast and slow. (Farrar, Straus & Giroux, 2011).

14 Ghosh, A., Rho, Y., McIntosh, A. R., Kotter, R. & Jirsa, V. K. Cortical network dynamics with time delays reveals functional connectivity in the resting brain. Cogn Neurodyn 2, 115-120, doi:10.1007/s11571-008-9044-2 (2008).

15 Deco, G., Jirsa, V., McIntosh, A. R., Sporns, O. & Kotter, R. Key role of coupling, delay, and noise in resting brain fluctuations. Proceedings of the National Academy of Sciences of the United States of America 106, 10302-10307, doi:0901831106 [pii] 10.1073/pnas.0901831106 (2009).

16 Cabral, J., Kringelbach, M. L. & Deco, G. Exploring the network dynamics underlying brain activity during rest. Prog Neurobiol 114, 102-131, doi:10.1016/j.pneurobio.2013.12.005 (2014).

17 Schirner, M., Rothmeier, S., Jirsa, V. K., McIntosh, A. R. & Ritter, P. An automated pipeline for constructing personalized virtual brains from multimodal neuroimaging data. NeuroImage, doi:10.1016/j.neuroimage.2015.03.055 (2015).

18 Schirner, M., McIntosh, A. R., Jirsa, V., Deco, G. & Ritter, P. Bridging multiple scales in the human brain using computational modeling. bioRxiv, http://dx.doi.org/10.1101/085548 (2016).

19 Deco, G., Jirsa, V. K. & McIntosh, A. R. Emerging concepts for the dynamical organization of resting-state activity in the brain. Nature reviews. Neuroscience 12, 43-56, doi:10.1038/nrn2961 (2011).

20 Deco, G., Jirsa, V. K. & McIntosh, A. R. Resting brains never rest: computational insights into potential cognitive architectures. Trends in neurosciences 36, 268-274, doi:10.1016/j.tins.2013.03.001 (2013).

21 Damoiseaux, J. S. et al. Consistent resting-state networks across healthy subjects. Proceedings of the National Academy of Sciences of the United States of America 103, 13848-13853 (2006).

22 Cabral, J. et al. Exploring mechanisms of spontaneous functional connectivity in MEG: How delayed network interactions lead to structured amplitude envelopes of band-pass filtered oscillations. Neuroimage 90, 423-435, doi:10.1016/j.neuroimage.2013.11.047 (2014).

23 Kuznetsov, Y. A. Elements of applied bifurcation theory. (Springer, 1998).

24 Freyer, F. et al. Biophysical mechanisms of multistability in resting-state cortical rhythms. The Journal of neuroscience: the official journal of the Society for Neuroscience 31, 6353-6361, doi:10.1523/JNEUROSCI.6693-10.2011 (2011).

25 Freyer, F., Roberts, J. A., Ritter, P. & Breakspear, M. A canonical model of multistability and scale-invariance in biological systems. PLoS Comput Biol 8, e1002634, doi:10.1371/journal.pcbi.1002634 (2012).

26 Hansen, E. C., Battaglia, D., Spiegler, A., Deco, G. & Jirsa, V. K. Functional connectivity dynamics: modeling the switching behavior of the resting state. NeuroImage 105, 525-535, doi:10.1016/j.neuroimage.2014.11.001 (2015).

27 Deco, G. & Jirsa, V. K. Ongoing cortical activity at rest: criticality, multistability, and ghost attractors. The Journal of neuroscience: the official journal of the Society for Neuroscience 32, 3366-3375, doi:10.1523/JNEUROSCI.2523-11.2012 (2012).

28 Deco, G. et al. Resting-state functional connectivity emerges from structurally and dynamically shaped slow linear fluctuations. The Journal of neuroscience: the official journal of the Society for Neuroscience 33, 11239-11252, doi:10.1523/JNEUROSCI.1091-13.2013 (2013).

29 Wildie, M. & Shanahan, M. Metastability and chimera states in modular delay and pulse-coupled oscillator networks. Chaos 22, 043131, doi:10.1063/1.4766592 (2012).

30 Kelso, J. A. S. Dynamic Patterns: The Self-Organization of Brain and behavior., (MIT Press, 1995).

31 Llinas, R. R. The intrinsic electrophysiological properties of mammalian neurons: insights into central nervous system function. Science 242, 1654-1664 (1988).

32 Varela, F., Lachaux, J. P., Rodriguez, E. & Martinerie, J. The brainweb: phase synchronization and large-scale integration. Nature reviews. Neuroscience 2, 229-239, doi:10.1038/35067550 (2001).

33 Haken, H. Information and Self-Organization. A macroscopic approach to Complex Systems. (Springer 1988).

34 Kelso, J. A. S. & Tognoli, E. in Neurodynamics of Cognition and Consciousness (eds R. Kozma & L. Perlovsky) 39-60 (Springer, 2007).

35 Rabinovich, M., Huerta, R., Varona, P. & Afraimovich, V. Transient Cognitive Dynamics, Metastability, and Decision Making. PLoS computational biology 4, e1000072 (2008).

36 Rabinovich, M. I., Huerta, R. & Laurent. Transient Dynamics for Neural Processing. Science 321, 48-50 (2008).

37 Huys, R., Perdikis, D. & Jirsa, V. K. Functional architectures and structured flows on manifolds: a dynamical framework for motor behavior. Psychol Rev 121, 302-336, doi:10.1037/a0037014 (2014).

38 Perdikis, D., Huys, R. & Jirsa, V. K. Time scale hierarchies in the functional organization of complex behaviors. PLoS computational biology 7, e1002198, doi:10.1371/journal.pcbi.1002198 (2011).

39 Beckmann, C. F., DeLuca, M., Devlin, J. T. & Smith, S. M. Investigations into resting-state connectivity using independent component analysis. Philos. Trans. R. Soc. Lond. B Biol. Sci. 360, 1001-1013 (2005).

40 Greicius, M. D., Krasnow, B., Reiss, A. L. & Menon, V. Functional connectivity in the resting brain: a network analysis of the default mode hypothesis. Proc. Natl Acad. Sci. USA 100, 253-258 (2003).

41 Mueller, S. et al. Individual variability in functional connectivity architecture of the human brain. Neuron 77, 586-595, doi:10.1016/j.neuron.2012.12.028 (2013).

42 Baker, A. P. et al. Fast transient networks in spontaneous human brain activity. eLife 3, e01867, doi:10.7554/eLife.01867 (2014).

43 Allen, E. A. et al. Tracking whole-brain connectivity dynamics in the resting state. Cerebral cortex 24, 663-676, doi:10.1093/cercor/bhs352 (2014).

44 Ghosh, A., Rho, Y., McIntosh, A. R., Kotter, R. & Jirsa, V. K. Noise during rest enables the exploration of the brain's dynamic repertoire. PLoS computational biology 4, e1000196, doi:10.1371/journal.pcbi.1000196 (2008).

45 Ponce-Alvarez, A. et al. Resting-state temporal synchronization networks emerge from connectivity topology and heterogeneity. PLoS computational biology 11, e1004100, doi:10.1371/journal.pcbi.1004100 (2015).

46 Gusnard, D. A. & Raichle, M. E. Searching for a baseline: functional imaging and the resting human brain. Nature Reviews Neuroscience 2, 685-694 (2001).

47 Addis, D. R., Wong, A. T. & Schacter, D. L. Remembering the past and imagining the future: common and distinct neural substrates during event construction and elaboration. Neuropsychologia 45, 1363-1377 (2007).

48 Botvinick, M., Nystrom, L. E., Fissell, K., Carter, C. S. & Cohen, J. D. Conflict monitoring versus selection-for-action in anterior cingulate cortex. Nature 402, 179-181, doi:10.1038/46035 (1999).

49 Goodkind, M. et al. Identification of a Common Neurobiological Substrate for Mental Illness. JAMA psychiatry 72, 305-315, doi:10.1001/jamapsychiatry.2014.2206 (2015).

50 Deco, G. & Kringelbach, M. L. Great Expectations: Using Whole-Brain Computational Connectomics for Understanding Neuropsychiatric Disorders. Neuron 84, 892-905 (2014).

51 Ritter, P., Schirner, M., McIntosh, A. R. & Jirsa, V. K. The virtual brain integrates computational modeling and multimodal neuroimaging. Brain connectivity 3, 121-145, doi:10.1089/brain.2012.0120 (2013).

52 Sanz Leon, P. et al. The Virtual Brain: a simulator of primate brain network dynamics. Frontiers in neuroinformatics 7, 10, doi:10.3389/fninf.2013.00010 (2013).

53 Ritter, P. & Villringer, A. Simultaneous EEG-fMRI. Neurosci.Biobehav.Rev. 30, 823-838 (2006).

54 Ritter, P., Becker, R., Freyer, F. & Villringer, A. in EEG-fMRI Physiology, Technique and Application (eds C. Mulert & L. Lemieux) Ch. 9, 153-171 (Springer, 2010).

55 Desikan, R. S. et al. An automated labeling system for subdividing the human cerebral cortex on MRI scans into gyral based regions of interest. NeuroImage 31, 968-980 (2006).

56 Tournier, J. D., Calamante, F., Gadian, D. G. & Connelly, A. Direct estimation of the fiber orientation density function from diffusion-weighted MRI data using spherical deconvolution. Neuroimage 23, 1176-1185, doi:10.1016/j.neuroimage.2004.07.037 (2004).

57 Kuramoto, Y. Chemical Oscillations, Waves, and Turbulence. Springer-Verlag, Berlin (1984).

58 Pikovsky, A., Rosenblum, M. & Kurths, J. Synchronization: a universal concept in nonlinear sciences. (Cambridge University Press, 2003).

59 Matthews, P. C. & Strogatz, S. H. Phase diagram for the collective behavior of limit-cycle oscillators. Phys. Rev. Lett. 65, 1701-1704 (1990).

60 Aronson, D. G., Ermentrout, G. B. & Kopell, N. Amplitude response of coupled oscillators. Physica D: Nonlinear Phenomena 41, 403-449 (1990).

61 Glerean, E., Salmi, J., Lahnakoski, J. M., Jaaskelainen, I. P. & Sams, M. Functional magnetic resonance imaging phase synchronization as a measure of dynamic functional connectivity. Brain connectivity 2, 91-101, doi:10.1089/brain.2011.0068 (2012).

62 Biswal, B., Yetkin, F. Z., Haughton, V. M. & Hyde, J. S. Functional connectivity in the motor cortex of resting human brain using echo-planar MRI. Magn.Reson.Med. 34, 537-541 (1995).

63 Buckner, R. L. et al. Cortical hubs revealed by intrinsic functional connectivity: mapping, assessment of stability, and relation to Alzheimer's disease. J Neurosci 29, 1860-1873, doi:10.1523/JNEUROSCI.5062-08.2009 (2009).

64 Achard, S., Salvador, R., Whitcher, B., Suckling, J. & Bullmore, E. A resilient, low-frequency, small-world human brain functional network with highly connected association cortical hubs. The Journal of neuroscience: the official journal of the Society for Neuroscience 26, 63-72, doi:26/1/63 [pii] 10.1523/JNEUROSCI.3874-05.2006 (2006).

